# Sponge contribution to the silicon cycle of a diatom-rich shallow bay

**DOI:** 10.1101/2021.10.20.465147

**Authors:** María López-Acosta, Manuel Maldonado, Jacques Grall, Axel Ehrhold, Cèlia Sitjà, Cristina Galobart, Aude Leynaert

## Abstract

In coastal systems, planktonic and benthic silicifiers compete for the pool of dissolved silicon, a nutrient required to make their skeletons. The contribution of planktonic diatoms to the cycling of silicon in coastal systems is often well characterized, while that of benthic silicifiers such as sponges has rarely been quantified. Herein, silicon fluxes and stocks are quantified for the sponge fauna in the benthic communities of the Bay of Brest (France). A total of 45 siliceous sponge species living in the Bay account for a silicon standing stock of 1215 tons, while that of diatoms is only 27 tons. The silicon reservoir accumulated as sponge skeletons in the superficial sediments of the Bay rises to 1775 tons, while that of diatom skeletons is only 248 tons. These comparatively large stocks of sponge silicon were estimated to cycle two orders of magnitude slower than the diatom stocks. Sponge silicon stocks need years to decades to be renewed, while diatom turnover lasts only days. Although the sponge monitoring over the last 6 years indicates no major changes of the sponge stocks, our results do not allow to conclude if the silicon sponge budget of the Bay is at steady state, and potential scenarios are discussed. The findings buttress the idea that sponges and diatoms play contrasting roles in the marine silicon cycle. The budgets of these silicon major users need to be integrated and their connections revealed, if we aim to reach a full understanding of the silicon cycling in coastal ecosystems.

## Introduction

There is great interest in understanding the cycling of silicon (Si) in marine environments because this nutrient is key to the functioning of marine ecosystems. In coastal oceans, Si is responsible for sustaining a large proportion of primary productivity and many of the food webs that ultimately sustain fish and human populations (Kristiansen and Hoell 2002; Ragueneau et al. 2006). Shortage of Si availability in coastal areas frequently reflects situations of ecosystem disequilibrium and proliferation of harmful algal blooms (Davidson et al. 2014; Glibert and Burford 2017; Thorel et al. 2017). Thus, a thorough understanding of the cycling of Si in coastal marine environments is critical for effective ecosystem management.

A substantial part of the Si cycling in the marine environment occurs through a variety of micro- and macro-organisms, the silicifiers, which require Si to build their siliceous skeletons (DeMaster 2003; Tréguer et al. 2021). To date, our understanding of the biogeochemical cycling of Si in the marine environment is based predominantly on the role of diatoms, microscopic unicellular eukaryotic algae which are the most abundant silicifiers in the global ocean (Malviya et al. 2016; Tréguer et al. 2021). Other silicifiers, such as siliceous Rhizaria and sponges, have received less attention and, consequently, their role is largely dismissed in most budgets of the marine Si cycle. To quantify the cycle contribution of siliceous sponges is particularly complicated given their benthic nature and heterogeneous distribution across the depths of the world’s oceans. However, there is growing evidence that sponges are important contributors to the Si cycle in terms of Si standing stocks and reservoirs (Maldonado et al. 2010, 2019).

In the present study, the Bay of Brest (France) was monitored to assess the relative contribution of siliceous sponges to the Si budget of this emblematic coastal system. This Bay has been the subject of numerous ecological, biogeochemical, and physical studies and is currently one of the best-studied coastal ecosystems in Europe, in terms of both structure and ecosystem functioning (Le Pape et al. 1996; Del Amo et al. 1997; Chauvaud et al. 2000; Ragueneau et al. 2018). In this ecosystem, diatoms dominate the annual pelagic primary production (Quéguiner and Tréguer 1984; Del Amo et al. 1997). Other planktonic silicifiers such as silicoflagellates, polycystines, and phaeodarians are rarely recorded in the monthly surveys of the planktonic community of the Bay, and when found, they are low abundant (https://www.phytobs.fr/). Silicic acid (dSi) concentrations vary over the year cycle from below 1 μM in spring and early summer up to 15-20 μM in late autumn and winter. Therefore, in this ecosystem diatom’s activity is limited by dSi principally during spring and early summer (Ragueneau et al. 2002), but sponges are limited all year round, as these organisms need dSi concentrations of 100-200 μM to reach their maximum speed of Si consumption (K_M_ = 30-100 μM Si; Reincke and Barthel 1997; Maldonado et al. 2011, 2020; López-Acosta et al. 2016, 2018).

In this system, both dredging and diving research activities have identified important populations of siliceous sponges (Jean 1994; López-Acosta et al. 2018), and a previous study firstly estimated that the sponge silica production represents ca. 8% of the net annual silica production in the Bay (López-Acosta et al. 2018). Nevertheless, the large amounts of Si within the sponge bodies (i.e., the Si standing stock) remain unquantified, as well as the fate of such Si once the sponges die. Herein, we are estimating these parameters and providing the most complete cycle of Si through sponges for this coastal ecosystem. The findings are discussed within the regional Si budget of the Bay of Brest previously published by Ragueneau et al. (2005), which considered only the contribution of planktonic diatoms.

## Materials

### Study area

The Bay of Brest (NW France) is a semi-enclosed marine ecosystem of about 130 km^2^ (harbors and estuaries not included) that is connected to the Atlantic Ocean through a narrow (1.8 km wide) and deep (45 m) strait. The Bay is a shallow system, with a maximum depth of 40 m and a mean depth of 8 m. It is a macro-tidal (maximum tidal amplitude = 8 m) system that receives high nutrient loadings mainly from two small rivers (Fig. 1a). During all the year cycle, the tidal and wind currents together with the shallowness of the Bay make nutrient concentration to remain relatively homogeneous in the water column (Delmas and Tréguer 1985; Salomon and Breton 1991; Le Pape et al. 1996).

**Fig. 1.**
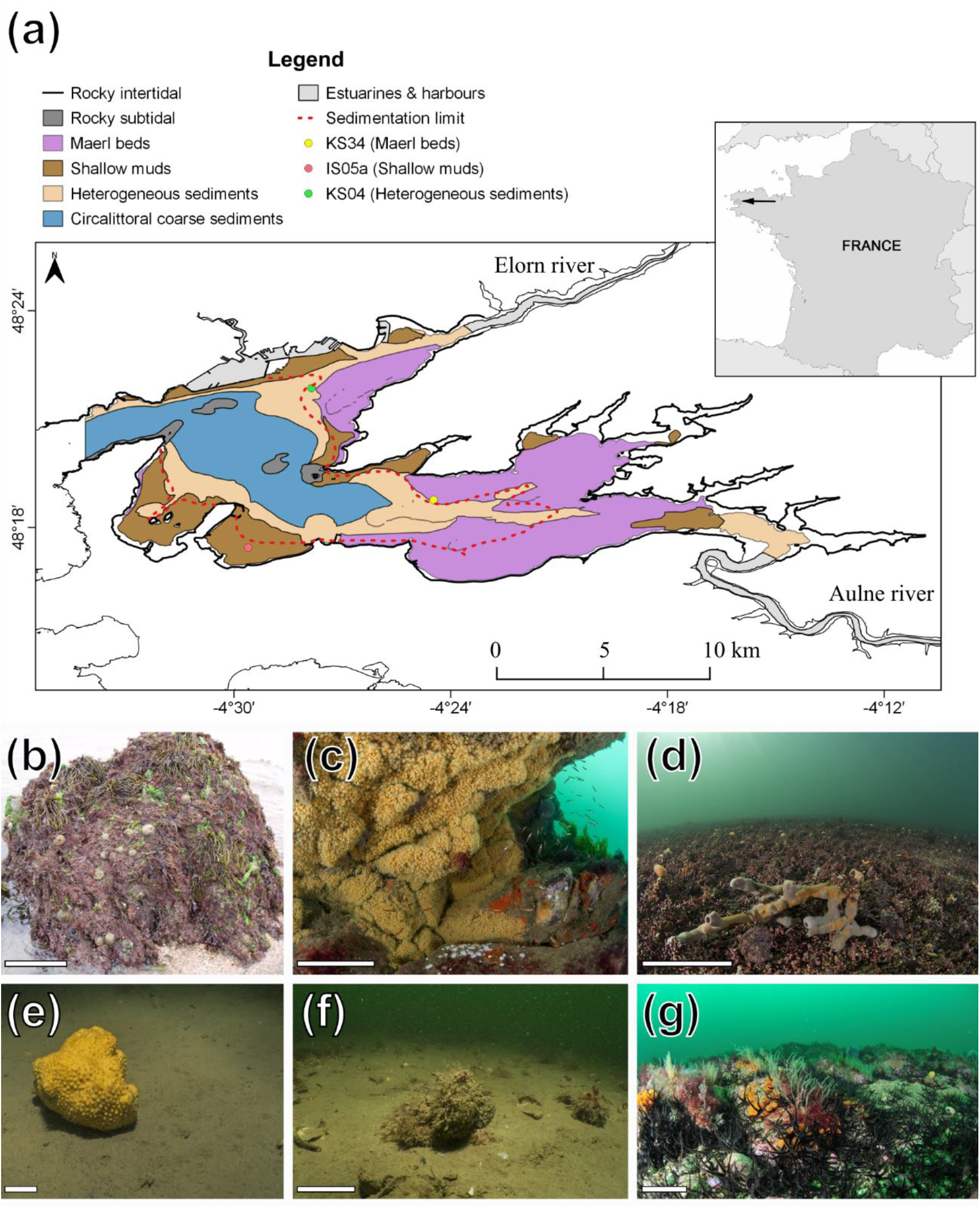
**(a)** Map of the Bay of Brest (France), showing the six major habitats defined for this study according to their depth, substrate type, and sponge fauna. Estuaries and harbors, depicted in grey in the map, have not been considered in the study. Red dashed line indicates the limit of sedimentation in the Bay, under which the sedimentation rate is negligible. Circles indicate the geographical location of the examined cores. (**b-g**) General view of the habitats of the Bay: (**b**) rocky intertidal, (**c**) rocky subtidal, (**d**) maerl beds, (**e**) shallow muds, (**f**) heterogeneous sediments, and (**g**) circalittoral coarse sediments. Scale bars indicate 10 cm.

The Bay hosts abundant and diverse benthic communities, developed in a mixture of hard and soft substrates (Grall and Glémarec 1997). The benthic ecosystem of the Bay consist of six major habitats, according to depth, nature of substrate, and biota (Hily et al. 1992; Gregoire et al. 2016): 1) the rocky intertidal coastline that remains emerged during spring low tides; 2) the rocky subtidal bottoms, down to 20 m depth, that mostly surround the islets of the Bay; 3) the maerl beds, soft bottoms with dense assemblages of *Lithophyllum* spp. down to 15 m depth; 4) the shallow mud bottoms down to 10 m depth, mostly in the peripheral zones of the Bay; 5) the heterogeneous sediments from 5 to 25 m depth which consist of a mix of mud, calcareous detritus, and gravel bottoms; and 6) the circalittoral coarse sediments, consisting of gravel and pebble bottoms located mostly in the central, deepest zones (20 to 40 m) of the Bay (Fig. 1a-g). Using QGIS software, version 3.10.2 (QGIS Development Team 2020), the benthic habitats of the Bay were delimited and their areas were calculated (Fig. 1a; Table 1). This resulted in the most updated map of the benthic habitats of the Bay of Brest, which includes the up-to-date information of the bottom communities of the Bay from the annual monitoring conducted by the Marine Observatory of the European Institute for Marine Studies (Derrien-Courtel et al. 2019).

**Table 1.** Features of the benthic habitats of the Bay of Brest (France), including bottom area (106 m^2^), relative contribution (%), sampled area (m^2^), mean (±SD) depth (m), and richness of siliceous species of each habitat and the total Bay. RI, rocky intertidal; RS, rocky subtidal; MB, maerl beds; SM, shallow muds; HS, heterogeneous sediments; CS, circalittoral coarse sediments.

| Habitat | Bottom area<br>(10<sup>6</sup> m<sup>2</sup>) | Sampled area<br>(m<sup>2</sup>) | Depth<br>(m) | Si species<br>richness |
| --- | --- | --- | --- | --- |
| RI | 1.98 (1.5%) | 23 | 0.0 (± 0.0) | 4 |
| RS | 2.64 (2.0%) | 31 | 12.4 (± 4.0) | 31 |
| MB | 45.84 (34.4%) | 40 | 5.4 (± 1.6) | 19 |
| SM | 22.19 (16.7%) | 17 | 5.1 (± 3.0) | 2 |
| HS | 31.79 (23.9%) | 36 | 11.4 (± 3.7) | 16 |
| CS | 28.69 (21.6%) | 23 | 25.6 (± 4.5) | 32 |
| Total | 133.13 (100%) | 170 | 9.9 (± 8.1) | 45 |

### Silicon standing stock in sponge communities

To estimate the Si standing stock of the sponge communities within the Bay of Brest, quantitative surveys were conducted across the six benthic habitats. The sampling effort carried out within a given habitat depended on both its relative surface to the total Bay extension and the relative internal variation (Eberhardt 1978). Three sampling techniques were used for the different habitats: (1) the rocky intertidal was sampled by using 23 random quadrats (1×1 m) at low tides; (2) the rocky subtidal, the maerl beds, the shallow mud, and the heterogeneous sediments above 20 m depth were sampled by scuba diving using 119 random 1×1 m quadrats; (3) the heterogeneous sediment and circalittoral coarse sediment located below 20 m depth were sampled using an epibenthic trawl over small transects (1 m wide × 5–10 m long, trawl length precision = ± 1m; n= 28). Each sponge found within the quadrats or in the trawls was counted, taxonomically identified to the species level, and measured for volume and silica content (i.e., biogenic silica, bSi). To determine the sponge volume, the body shape of each individual was approximated to one or a sum of several geometric figures (e.g., sphere, cylinder, cone, rectangular plate, etc.) and the linear parameters needed to calculate the volume (length, width, and/or diameter) were measured in situ using rulers (Maldonado et al. 2010). Counts and volume values in each habitat were normalized to m^2^. Non-siliceous sponge species were taxonomically identified but not considered in the calculations.

The relationship between the mean siliceous sponge abundance (i.e., number of individuals) and biomass (i.e., volume) in each habitat was normalized per m^2^ and analyzed by Spearman rank correlation. Between-habitat differences in the sponge abundance and biomass per m^2^ were examined by Kruskal-Wallis analysis when data did not meet normality and/or equal variance, following Shapiro-Wilk and Brown-Forsythe tests, respectively. Post-hoc pairwise multiple comparisons between groups were conducted using the non-parametric Dunn’s test.

To estimate sponge silica content in a given sponge volume, a plastic cylinder of known volume was filled with sponge tissue (n = 3 to 5 for each species, depending on availability), applying minimum compression. Sponge tissue samples were then dried at 60°C to constant dry weight (g), and desilicified by immersion in 5% hydrofluoric acid solution for 5 hours, rinsed in distilled water three times for 5 minutes, and dried again at 60°C to constant dry weight (Maldonado et al. 2010). Silica content per unit of sponge volume (mg bSi mL^−1^ sponge) was calculated as the difference in weight before and after desilicification and multiplied by a factor of 0.8. Such a factor was applied to remove overestimates of the skeletal biogenic silica due to tiny sand grains (lithogenic silica) that might be embedded into the sponge tissues (Maldonado et al. 2010).

Mean skeletal content (mg bSi mL^−1^ sponge) of each species was subsequently used to estimate Si standing stock in the sponge communities per unit of bottom area in the six habitats of the Bay, and that at the ecosystem level. Between-habitat differences in mean sponge Si standing stock per m^2^ were examined by a Kruskal-Wallis analysis, followed by pairwise Dunn’s tests to identify significant differences between groups. Finally, the relationship between skeletal content and sponge abundance per m^2^ and that between skeletal content and sponge biomass per m^2^ in the sponge communities of the Bay were also examined using regression analysis.

### Dissolved silicon consumption by sponge communities

The annual consumption of dSi (and consequently biogenic silica production) by the sponge communities of the Bay was estimated by using the dSi consumption kinetic models developed elsewhere (López-Acosta et al. 2016, 2018) for the four dominant sponge species at the Bay (*Haliclona simulans*, *Hymeniacidon perlevis*, *Tethya citrina*, and *Suberites ficus*). By knowing the monthly average dSi availability in the bottom waters of the Bay over the last decade (2012-2021; Table S1) and the consumption kinetics of each of the four dominant species at the Bay, the average (±SD) annual dSi consumption rate by these species was estimated. This average was then extrapolated to the rest of the sponge species in the Bay to estimate dSi consumption by habitat and for the entire Bay, according to the previously estimated species sponge biomass per habitat (mL sponge m^−2^) and habitat bottom area (m^2^).

### Sponge biogenic silica in superficial sediments

The amount of sponge Si in the sediments was determined by analyzing superficial sediments samples (defined herein as the upper-centimeter layer of sediment) from three stations from the shallow plateaus of the Bay of Brest (Fig. 1a). In the Bay, there are two major depositional environments with contrasting features: 1) the shallow terraces or plateaus, up to 10 m depth, where the fine sediment that regularly arrives through the rivers’ discharges is homogeneously deposited, and 2) the deepest bottoms, up to 40 m depth, where the strong tidal currents prevent the fine sediment from being deposited on the bottom (Salomon and Breton 1991; Gregoire et al. 2017; Lambert et al. 2017). The latter are located mainly in the central zone of the Bay and the two paleo-channels that cross each of the basins of the Bay from the rivers’ mouths to the central zone, where they converge (see Figure 1 in Gregoire et al. 2017). This depositional configuration was originated by the action of successive sea-level low stands and it has been preserved over the last millennia by the effect of tidal currents in deep-water areas, which keep the inherited shape by preventing the deposition of the fine sediment supplied by rivers (Gregoire et al. 2016, 2017). As a consequence of this hydrodynamic regime, the fine sediment does not settle on the deepest areas of the Bay, the sediments of which mainly consist of coarse particles such as gross fragments of shells and small pebbles (Gregoire et al. 2016). On the contrary, the fine sediment settles all over the shallow plateaus, which are less affected by the tidal currents. Hence, the shallow plateaus account for virtually the total area of sediment deposition in the Bay bottoms (Ehrhold et al. 2016). These shallow plateaus, which account for about half of the Bay surface, are mainly covered by maerl beds, shallow muds and partially by heterogeneous sediments (Fig. 1a; Table 2).

**Table 2.**
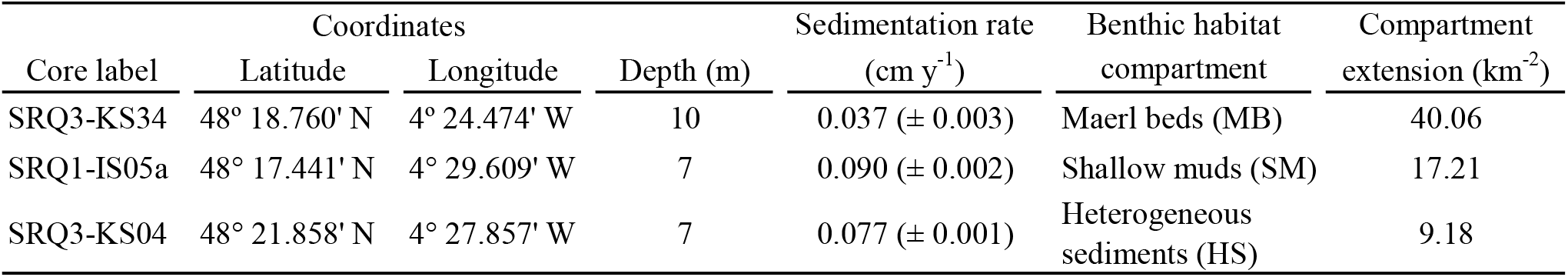
Summary of core features including core label, coordinates, depth (m), average (±SD) sedimentation rate (cm y^−1^), the benthic habitat compartment they represent, and the extension of the benthic habitat over the sedimentation limit of the Bay (km^2^). Present-day sedimentation rates for the superficial sediments of each core were estimated from ^14^C radiocarbon dating in Gregoire et al. (2017) and Ehrhold et al. (2021).

To estimate the amount of sponge silica in the superficial sediments of the Bay, one core was sampled at each benthic habitat represented at the shallow plateaus: core SRQ3-KS34 from maerl beds, core SRQ1-IS05 from shallow muds, and core SRQ3-KS04 from heterogeneous sediments (Fig. 1a, Table 2). Present-day sedimentation rates were estimated from radiocarbon dating of superficial sediment in the cores (Gregoire et al. 2017; Ehrhold et al. 2021). The bottoms of the deepest areas of the Bay, on which sedimentation rates are negligible (Ehrhold et al. 2016), were interpreted as null contributors to the sponge silica accumulation in the Bay (see below).

To quantify the sponge silica in the superficial sediment layer, one to three sediment subsamples of 10 mg each were collected from the upper 1-cm layer of each core and subsequently processed by light microscopy, following the methodology described in Maldonado et al. (2019). Briefly, sediment subsamples were transferred to test tubes, boiled in 37% hydrochloric acid to remove calcareous materials, then in 69% nitric acid to complete digestion of organic matter, rinsed in distilled water, and sonicated for 15 minutes to minimize sediment aggregates. The sediment suspension was then pipetted out and dropped on a microscope glass slide to measure the volume of each skeletal piece. A total of 230 smear slides in which 48320 spicules, either entire or fragmented, were examined through contrast-phase compound microscopes. Using digital cameras and morphometric software (ToupView, ToupTek Photonics), the volume of each spicule or spicule’s fragment was estimated by approximating its shape to one or the sum of several geometric figures (Maldonado et al. 2019). Measured volumes of sponge silica were subsequently converted into Si mass, using the average density of 2.12 g mL^−1^ for sponge silica (Sandford 2003).

To determine the reservoir of sponge silica in the superficial sediment of the Bay, the obtained mass of sponge silica per gram of sediment in each core was extrapolated to the area of each benthic habitat over the sedimentation limit, delimited with a red dashed line in Figure 1 from the sedimentological data published in Ehrhold et al. (2016). It means that the sponge silica mass per gram of sediment from core SRQ3-KS34 was extrapolated to 40.06 km^2^, that of core SRQ1-IS05 to 17.21 km^2^, and that of core SRQ3-KS04 to 9.18 km^2^ (Table 2). Sponge Si deposition rate was estimated from the amount of sponge Si determined in the superficial sediments of each core and the sedimentation rate (Table 2). Sponge Si burial rate, i.e., the amount of sponge silica that is ultimately preserved in the sediments, was estimated from the rate of sponge Si deposited annually in the superficial sediments and the average percentage estimated for sponge Si preservation in sediments of marine continental shelves (i.e., 52.7 ± 29.8%; Maldonado et al., 2019). The contribution of sponges to the Si benthic efflux from the sediments of the Bay was calculated as the difference between the sponge Si deposition and burial rates.

### Sponge silica budget in a coastal ecosystem: the Bay of Brest

The quantified stocks and fluxes of sponge Si were used to build a sponge Si budget for the Bay of Brest. These results were discussed in the context of previous studies and compared with those reported in the literature for the community of planktonic diatoms in the Bay.

## Results

### General features of the sponge assemblages

The survey of the sponge fauna at the Bay of Brest revealed a total of 53 sponge species, most of them belonging to the class Demospongiae (n=51), and only 2 species belonging to the class Calcarea (Table S2). Species from classes Hexactinellida and Homoscleromorpha were not found. Forty-five (85%) of the 53 identified species had siliceous skeletons. The other 8 were non-siliceous sponge species with a skeleton made of either calcium carbonate (n=2) or skeletal protein (n=6). Non-siliceous sponges were not considered in the following quantifications of sponge abundance and biomass.

Sampling yielded a total of 1807 sponge individuals for which volume was determined (Table 1). At the Bay level, the siliceous sponge fauna averaged 11.0 ± 11.1 individuals m^−2^ and a biomass of 0.31 ± 0.35 L of living sponge tissue m^−2^. The large variation associated (i.e., standard deviation, SD) to the sponge abundance and biomass normalized per m^2^ is derived from the patchy spatial distribution of the sponges at both intra- and inter-habitat level. In contrast, the standard error (SE) of the mean (=SD/√N) is low (SE of sponge abundance = 0.85; SE of sponge biomass = 0.03), indicating that mean values are accurate. A Spearman’s correlation involving the six habitats revealed a strong positive linear relationship between the mean abundance and biomass of siliceous sponges per m^2^ of sampled bottom (N = 6, ρ = 0.886, *p* = 0.033). This relationship informs that body size is more or less uniformly distributed across species and specimens of the different habitats.

At the habitat level, there were large between-habitat differences in both sponge abundance and biomass (Fig. 2). The total number of siliceous sponges per m^2^ significantly differs between habitats (H= 89.059, df= 5, *p*< 0.001). Mean siliceous sponge abundance per m^2^ in the benthic communities of the maerl beds and rocky subtidal habitats (24.6 ± 26.2 and 16.4 ± 9.3 individuals m^−2^, respectively) were significantly higher than that in the other habitats (Fig. 2a). Biomass of siliceous sponges (L m^−2^) also differed significantly between habitats (H= 78.324, df= 5, *p*< 0.001). A posteriori pairwise comparison revealed that mean sponge biomass per m^2^ at the rocky subtidal habitat, which showed the highest sponge biomass per sampled bottom (3.8 ± 9.7 L m^−2^), was significantly higher than that at the heterogeneous sediments (0.16 ± 0.38 L m^−2^) and the rocky intertidal (0.12 ± 0.13 L m^−2^) habitats. Mean sponge biomass per m^2^ at maerl beds (0.31 ± 0.32 L m^−2^), which ranked second, did not differ significantly from that at the rocky subtidal, either at the heterogeneous sediments and rocky intertidal habitat. Mean sponge biomass per m^2^ at shallow muds and circalittoral coarse sediments were similar to each other and significantly lower than those at the rest of habitats in the Bay (Fig. 2b-c).

**Fig. 2.**
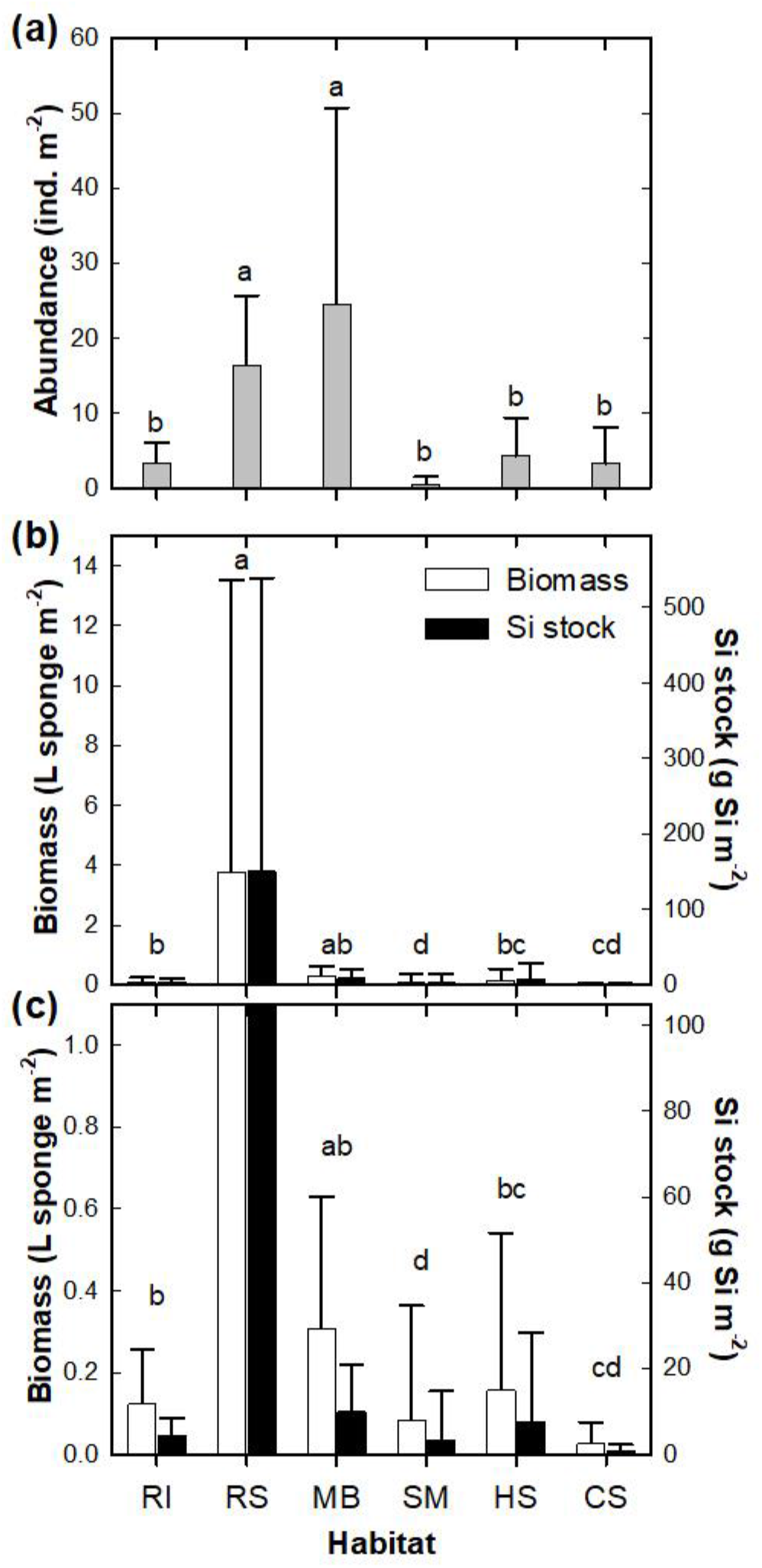
Summary of (a) average (±SD) abundance (individuals m^−2^), (b-c) biomass (L sponge m^−2^; white bars), and silicon (Si) standing stock (g Si m^−2^; black bars) in the siliceous sponge fauna of the habitats of the Bay of Brest (France). Significant between-habitat differences of (**a**) abundance and (**b**) biomass and Si stock of the sponge fauna are indicated with different letters according to the results of Kruskal-Wallis analysis and the a posteriori pairwise Dunn’s tests. (**c**) This graph makes visible the contribution of the habitats with sponge biomass and Si standing stock records lower than 1 L sponge m^−2^ and 100 g Si m^−2^. Habitats abbreviations are RI for rocky intertidal, RS for rocky subtidal, MB for maerl beds, SM for shallow muds, HS for heterogeneous sediments, and CS for circalittoral coarse sediments.

### Silicon standing stock in the sponge communities

The importance of the siliceous skeleton content per mass unit of sponge tissue varied largely between sponge species (Table 3). It ranged from 29.8 ± 2.9 to 145.6 ± 14.2 mg bSi per mL of living sponge tissue (Table 3), accounting for 19.6% to 63.5% of the specific dry weight (DW). The average skeletal content estimated for the siliceous sponge fauna of the Bay of Brest was 59.6 ± 27.1 mg bSi mL^−1^ (45.8 ± 10.6% bSi/DW). From these figures, it is estimated that the Si standing stock in the sponge fauna of the Bay of Brest is 12.3 ± 14.1 g Si m^−2^. Nevertheless, there are large between-habitat differences in the sponge Si standing stock per m^2^ (Fig. 2b-c). The highest mean Si standing stock occurred in the rocky subtidal habitat (151.1 ± 387.6 g Si m^−2^). At this habitat, large specimens of *Cliona celata* were common (mean abundance= 0.5 ± 0.7 individual m^−2^, mean biomass= 7.4 ± 13.5 L individual^−1^). The large specimens of this species, which is moderately silicified (85.6 ± 7.2 mg bSi mL^−1^; Table 3), are responsible for the 89.2% of the total biomass and Si standing stock of the rocky subtidal habitat. The maerl beds ranked second in Si standing stock, with an average of 10.0 ± 10.7 g Si m^−2^ (Fig. 2b-c). In this habitat there is large abundance of sponges (24.6 ± 26.2 individuals m^−2^) with body size ranging from small to medium (mean biomass= 12.4 ± 37.1 mL individual^−1^). Four of the 19 siliceous sponge species identified at this habitat accounted for 84% of the total sponge Si standing stock (*Haliclona simulans*, *Hymeniacidon perlevis*, *Tethya citrina*, and *Suberites ficus*). A Kruskal-Wallis analysis confirmed that the between-habitat differences in sponge Si standing stock per m^2^ are statistically significant (H = 71.701, df = 5, *p*<0.001). The pairwise comparison of mean Si standing stock revealed the same pattern of between-habitat differences than that obtained in the pairwise comparison of mean sponge biomass (Fig. 2b-c). Further analysis indicated a significant linear relationship (n= 170, R^2^= 0.999, *p*<0.0001) between sponge biomass and Si standing stock per m^2^ across habitats (Fig. 3).

**Table 3.**
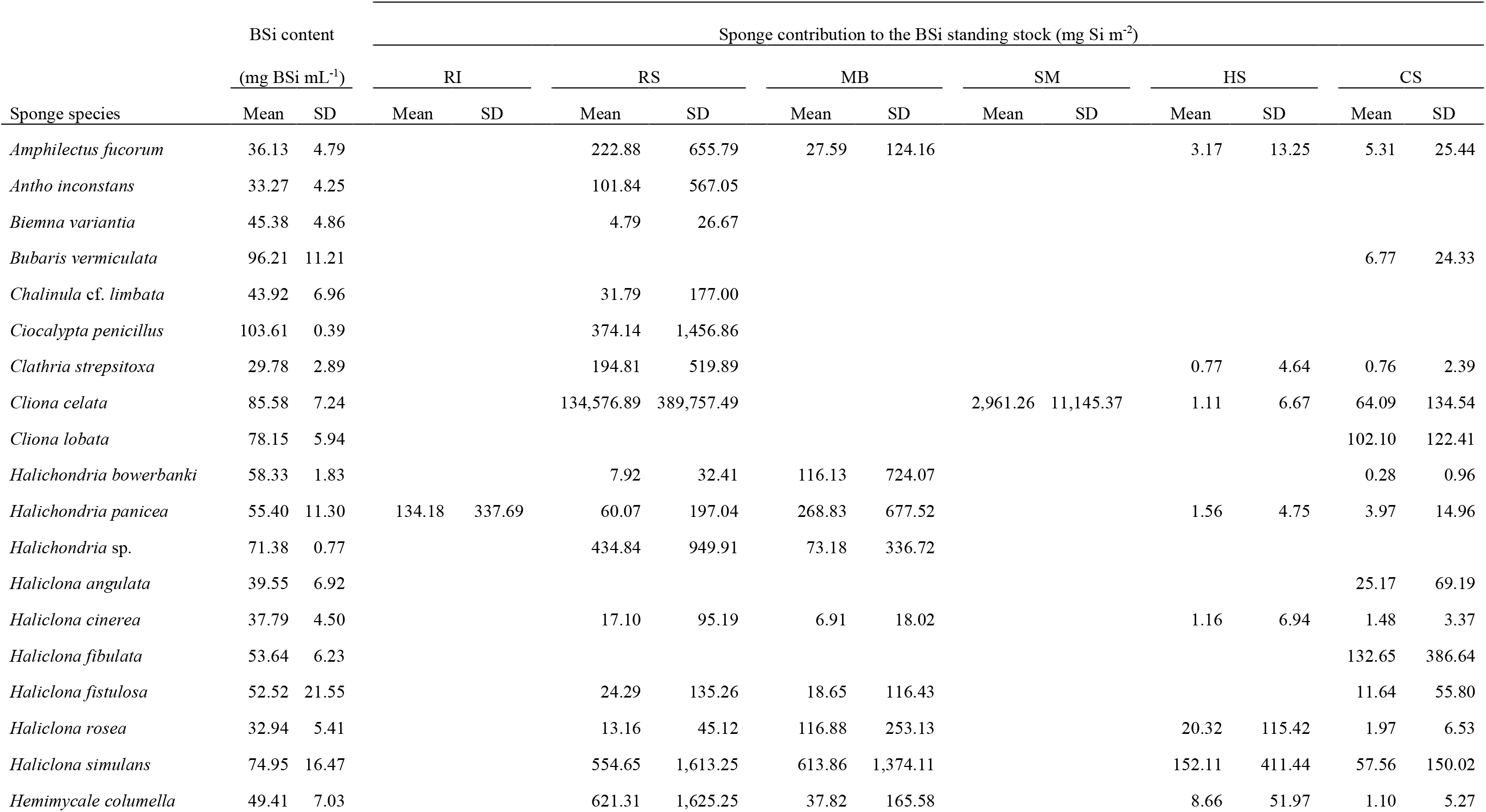
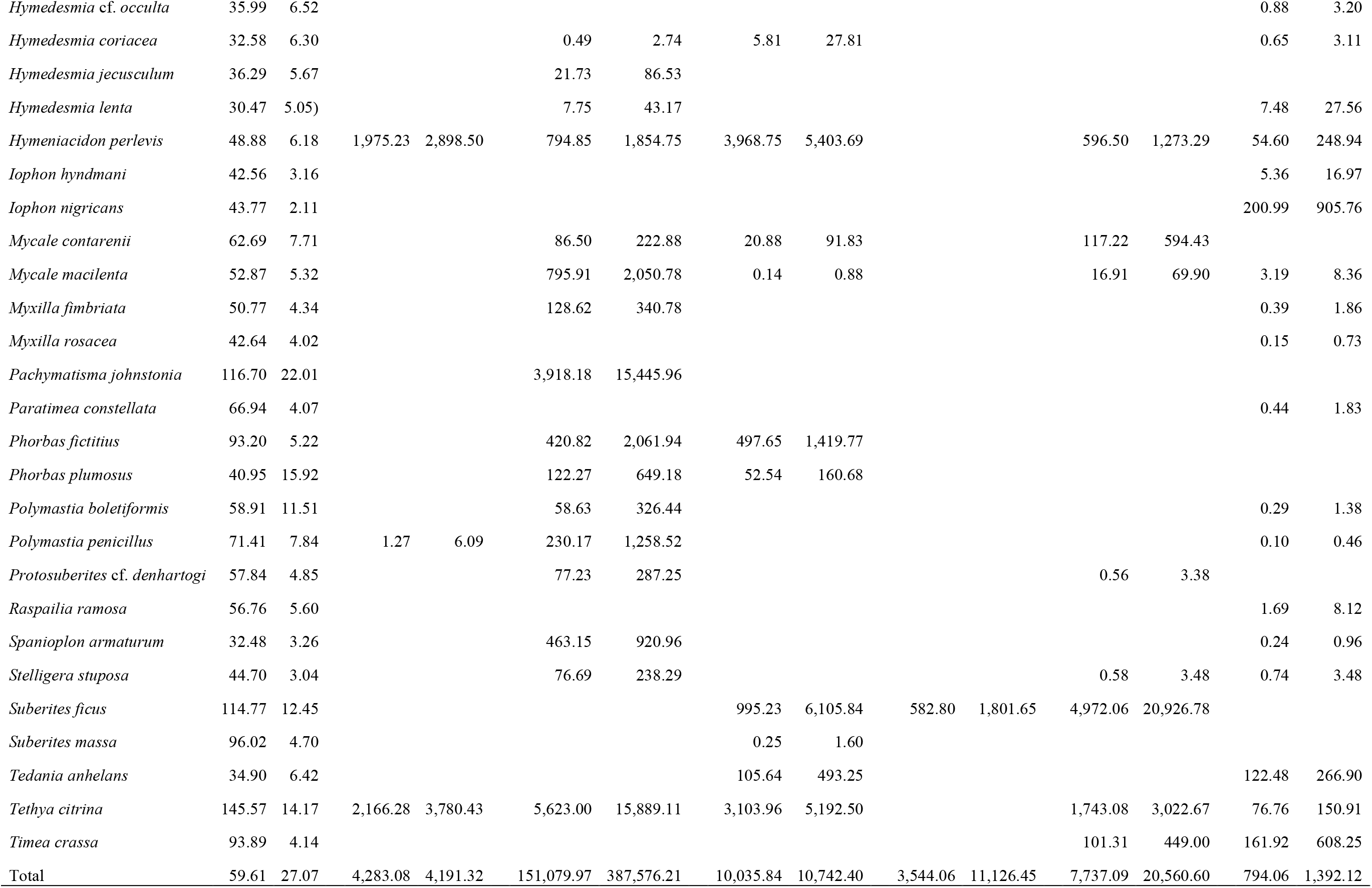
Summary of biogenic silica (bSi) standing stock in the sponge assemblages of the Bay of Brest (France). Average (±SD) silica content per unit of volume of living sponge tissue (mg bSi mL^−1^) of each siliceous sponge species and average (±SD) sponge Si stock per square meter (mg Si m^−2^) in each habitat (RI, rocky intertidal; RS, rocky subtidal; MB, maerl beds; SM, shallow mud; HS, heterogeneous sediments; CS, circalittoral coarse sediments) and for the total ecosystem of the Bay of Brest.

**Fig. 3.**
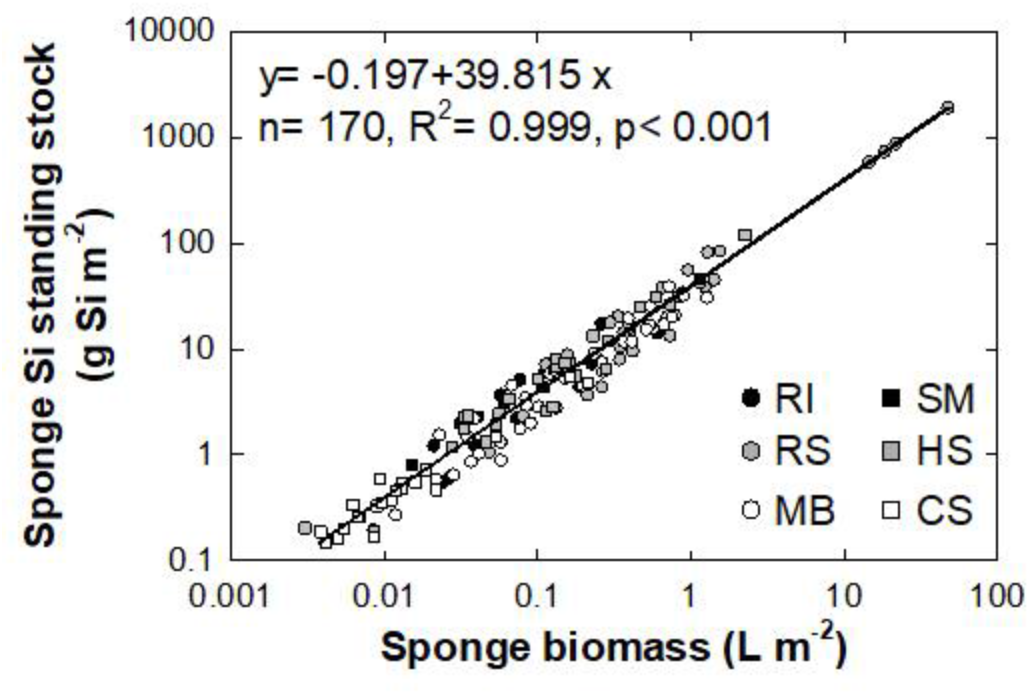
Relationship between silicon (Si) standing stock (g Si m^−2^) and biomass (L m^−2^) of siliceous sponges per m^2^ of sampled bottom at the Bay of Brest (France). Symbols represent the location of the data (i.e., habitat; abbreviations mean: RI, rocky intertidal; RS, rocky subtidal; MB, maerl beds; SM, shallow muds; HS, heterogeneous sediments; CS, circalittoral coarse sediments). Sponge Si stock and biomass are shown in logarithmic scale.

By integrating the average Si content of the sponges across the bottom area of each habitat, a total Si standing stock of 1215 ± 1876 × 10^3^ kg Si (43.3 ± 66.8 × 10^6^ mol Si) in the sponge communities of the Bay of Brest was estimated. The small-scale patchiness in the spatial distribution of the sponges within a habitat causes some sampling quadrats to contain many sponges while others contain very few or none. This effect, when propagated for the calculation of the global mean of the Bay, results in a large SD value. In addition, about 90% of the stock is accumulated in the sponge fauna of three habitats, which —in order of contribution— are the maerl beds, the rocky subtidal, and the heterogeneous sediments.

### Dissolved silicon consumption by sponge communities

The average monthly concentrations of dSi at the bottom waters of the Bay ranged from 2 to 13 μM (Table S1). At these nutrient concentrations, the most abundant species at the Bay (*H. perlevis*, 68.6 ± 85.1 mL m^−2^) consumed 7.2 ± 8.9 mmol Si m^−2^ y^−1^. Interestingly, the species *T. citrina* and *S. ficus*, which show comparatively lower biomass records in the Bay (24.5 ± 34.1 and 7.7 ± 13.3 mL m^−2^, respectively), had similar rates of Si consumption (7.5 ± 10.5 and 7.1 ± 21.0 mmol Si m^−2^ y^−1^, respectively). This is because their affinity for dSi is higher (affinity coefficient, V_max_/K_M_ = 6.9 × 10^−3^ and 4.4 × 10^−3^ μmol Si h^−1^ sponge-mL^−1^ Si-μM^−1^ for *T. citrina* and *S. ficus*, respectively, compared to 2.1 × 10^−3^ μmol Si h^−1^ sponge-mL^−1^ Si-μM^−1^ for *H. perlevis*).

Not surprisingly, most of the sponge Si consumption of the Bay occur in the rocky subtidal habitat and maerl beds, where sponges are very abundant and they show moderate to large biomass records (Fig. 2). The rocky subtidal habitat, which represents only 2.0% of the Bay bottom but hosts about 30% of the total sponge biomass in the Bay, shows an annual sponge Si consumption rate of 74.2 ± 141.4 × 10^3^ kg Si y^−1^ (2.6 ± 5.0 × 10^6^ mol Si y^−1^). The maerl beds, in which *H. perlevis, T. citrina*, and *S. ficus* are abundant in both terms of number of individuals and biomass, shows an annual Si consumption rate of 73.6 ± 112.7 × 10^3^ kg Si y^−1^ (2.6 ± 4.0 × 10^6^ mol Si y^−1^). Collectively, the assemblage of siliceous sponges in the Bay is estimated to consume annually 200.8 ± 372.1 × 10^3^ kg Si y^−1^ (7.2 ± 13.2 × 10^6^ mol Si y^−1^).

### Sponge bSi in sediments

The superficial sediments (i.e., the upper 1-cm layer) of cores SRQ3-KS34, SRQ1-IS05a, and SRQ3-KS04, contained respectively a total of 1.07 (±0.05), 1.30, and 0.32 million spicules or spicule fragments per g of sediment, with an average spicule content of 0.89 (±0.51) million spicules g^−1^ sediment for the set of studied cores (Table 4). In all cores, most spicules (98.3-99.1%) were recognized as megascleres, either entire or fragmented. The microscopic study of the superficial sediment in the cores showed only small between-area differences in the mass of sponge bSi, which ranged from 0.799 to 2.505 mg Si g^−1^ sediment (Table 4). The average content of sponge bSi in the sediments of the Bay of Brest was 1.565 (±0.866) mg Si g^−1^ sediment.

**Table 4.** Summary of the results obtained during the microscopic examination of the superficial sediments of the Bay of Brest. Average (±SD, when available) spicule counts (in millions per gram of sediment), contribution (%) of megascleres (spicules longer than 100 μm) and microscleres (spicules shorter than 100 μm), and mass of sponge silica per gram of sediment for each core are indicated, as well as for the set of study cores.

| Core label | Benthic habitat | N spicules<br>(N x 10<sup>6</sup> g<sup>-1</sup> sed.) | Spicule-type contribution (%) |  | Sponge Si in superficial sed.<br>(mg Si g<sup>-1</sup> sed.) |
| --- | --- | --- | --- | --- | --- |
|  |  |  | Megascleres | Microscleres |  |
| SRQ3-KS34 | Maerl beds (MB) | 1.07 (± 0.05) | 99.1 (± 0.3) | 0.9 (± 0.3) | 1.391 (± 0.241) |
| SRQ1-IS05a | Shallow muds (SM) | 1.30 | 99.0 | 1.0 | 2.505 |
| SRQ3-KS04 | Heterogeneous sediments (HS) | 0.32 | 98.3 | 1.7 | 0.799 |
| AVERAGE (±SD) |  | 0.89 (± 0.51) | 98.8 (± 0.4) | 1.2 (± 0.4) | 1.565 (± 0.866) |

Sponge Si deposition rates in the studied areas ranged from 9.4 to 64.4 × 10^3^ kg Si y^−1^, depending on the habitat (Table 5). When deposition rates were extrapolated to the total extension of the shallow plateaus (i.e., 66.45 km^2^), it resulted in a mean deposition rate of 108.7 ± 9.1 × 10^3^ kg Si y^−1^ (3.9 ± 0.3 × 10^6^ mol Si y^−1^). The sponge Si burial rate was then calculated from the deposition rate and the average preservation rate of sponge silica determined by Maldonado et al. (2019) for continental-shelf sediments. This approach yielded an average burial rate of sponge silica in the sediments of the shallow plateaus of 57.3 ± 18.2 × 10^3^ kg Si y^−1^ (2.0 ± 0.6 × 10^6^ mol Si y^−1^). The differences between deposition and burial rate is, in the long run, the sponge contribution to the benthic Si efflux from sediments, which was estimated to be 51.4 ± 17.7 × 10^3^ kg Si y^−1^ (1.8 ± 0.6 × 10^6^ mol Si y^−1^) for the shallow plateaus of the Bay (Table 5).

**Table 5.** Sponge silicon (Si) stock in the superficial sediments of each depositional environment of the Bay of Brest, along with the deposition, burial and benthic flux rate of sponge Si. Area (km^2^) to which the sponge Si data were extrapolated is indicated. These areas do not correspond to the total habitat extension but to the habitat area that is above the sedimentation limit of the Bay (indicated with a red dashed line in Fig. 1a) for cores representing the shallow depositional plateaus.

| Core label | Benthic habitat | Area | Sponge Si STOCK | Sponge Si FLUXES |  |  |
| --- | --- | --- | --- | --- | --- | --- |
|  |  | km<sup>2</sup> | reservoir<br>10<sup>3</sup> kg Si | deposition rate<br>10<sup>3</sup> kg Si y<sup>-1</sup> | burial rate<br>10<sup>3</sup> kg Si y<sup>-1</sup> | benthic flux<br>10<sup>3</sup> kg Si y<sup>-1</sup> |
| SRQ3-KS34 | Maerl beds (MB) | 40.06 | 937.93 | 34.91 | 18.39 | 16.52 |
| SRQ1-IS05a | Shallow muds (SM) | 17.21 | 715.41 | 64.39 | 33.92 | 30.47 |
| SRQ3-KS04 | Heterogeneous<br>sediments (HS) | 9.18 | 121.70 | 9.38 | 4.94 | 4.44 |
| TOTAL |  | 66.45 | 1775.04 | 108.68 | 57.25 | 51.43 |

According to the depositional environments of the Bay, the deep sediments were not considered in the calculations of the total sponge Si reservoir and sediment fluxes rates (see Methods). This is also supported by the sponge spatial distribution at the Bay, which shows that in the deepest areas sponges are particularly low abundant (3.2 ± 4.9 ind. m^−2^) and their contribution to the Si standing stock is significantly low (0.8 ± 1.4 g Si m^−2^; Fig. 2). By extrapolating the sponge silica content determined in the superficial sediments of the Bay (Table 5), a total sponge silica reservoir in the superficial (upper centimeter) sediment layer of the Bay of Brest of 1775 ± 162 × 10^3^ kg Si (63.2 ± 5.8 × 10^6^ mol Si) was estimated.

## Discussion

### Sponge fauna within the Bay of Brest

The survey of the sponge fauna of the Bay of Brest revealed that sponges are ubiquitous and widely present in most of the habitats (17-36 spp. per habitat; Table 1, Fig. 2, Table S2). Only two habitats, the shallow muds and the rocky intertidal, showed low sponge species richness (≤ 4 species per habitat; Table 1, Table S2). These two habitats, the former characterized by muddy bottoms and scanty hard substratum and the latter by periodic air exposures during low tides, hosted only a handful of species able to deal with those harsh conditions (Table S2).

The taxonomic composition and spatial distribution of the sponge fauna significantly differed between habitats (Table 1, Fig. 2). Spatial heterogeneity has been widely reported for sponge distributions across all oceans, with a variety of biological and physical reasons behind it (Hooper 2019). Among the habitats of the Bay, the rocky subtidal is the one with the highest sponge biomass per unit area (3.8 ± 9.7 L m^−2^; Fig. 2b), reaching values similar to those reported in some emblematic sponge aggregations in tropical and polar latitudes (Maldonado et al. 2017 and references therein). According to their extension, the maerl beds, which accounted for about one third of the total surface of the Bay (Table 1), hosted most of the sponge individuals and biomass of the Bay —79 ± 46% of the total number of sponges (1416 ± 1474 million of individuals) and about 44 ± 23% of the total sponge volume (31.9 ± 47.0 × 10^6^ L) estimated at the Bay of Brest. Thus, the maerl beds of the Bay serve as substrate to highly diverse and abundant sponge fauna, which along with other benthic organisms, make these bottoms real sponge and biodiversity hotspots, similar to what has been reported for maerl beds from other ocean regions (Sciberras et al. 2009; Ávila and Riosmena-Rodriguez 2011; Neill et al. 2015).

### Biogenic silica stock as sponge skeletons

Our results indicate that a substantial amount of the biogenic silica stock of the Bay of Brest is in the form of sponge silica, in both the living sponge fauna and the sediments of the Bay (Fig. 4). In the sponge communities, the Si standing stock within the skeleton of the sponges totaled 1215 ± 1876 × 10^3^ kg of Si. The siliceous skeleton accounted for 19.6-63.5% of the sponge dry weight, depending on the species (average value = 45.8 ± 10.6% bSi/DW = 59.6 ± 27.1 mg bSi mL^−1^; Table 3). Similar bSi content and between-species variability was also reported in a sponge-rich community of the Belizean Mesoamerican Barrier Reef (53.4 ± 6.1 mg bSi mL^−1^; Maldonado et al. 2010).

**Fig. 4.**
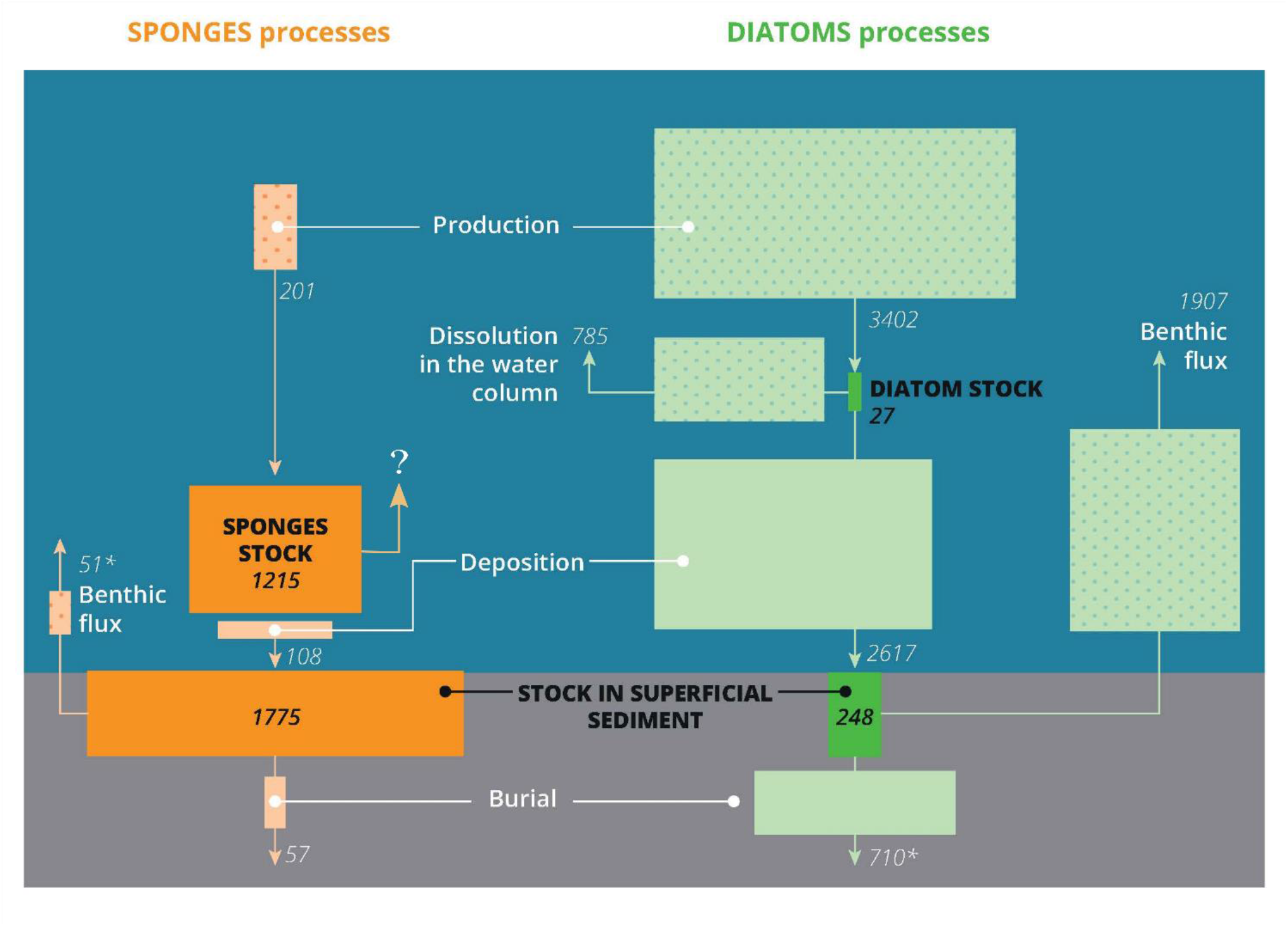
Scheme summarizing the silicon stocks and fluxes through the sponge and planktonic diatom communities of the Bay of Brest (France). Stocks and fluxes of silicon (Si) mediated by sponges are in orange, and those mediated by diatoms are in green. Stocks of biogenic silica are indicated in tons of Si. Fluxes of silicic acid, with a dotted pattern, and those of biogenic silica, lacking the dotted pattern, are indicated in tons of Si per year. The size of the boxes representing both stocks and fluxes is proportional to their rate. Fluxes of Si through diatoms are from Ragueneau et al. (2005), stock of biogenic silica in planktonic diatoms has been calculated from Beucher et al. (2004), and reservoir of diatom silica in superficial sediments are from Song and Ragueneau (2007); more details about calculations in the main text. Asterisks (*) refer to fluxes derived indirectly.

To date, only few studies have measured the amount of Si trapped in sponge communities of shallow-water ecosystems. The sponge Si standing stock per m^2^ in the Bay of Brest (12.3 ± 14.1 g Si m^−2^) is about 5 times higher than that found in the sublittoral population of the encrusting demosponge *Crambe crambe* at the Catalan coast of the NW Mediterranean (2.5 ± 2.7 g Si m^−2^; Maldonado et al. 2005) and, interestingly, similar to that measured in the sublittoral sponge-rich assemblages of the Belize section of the Mesoamerican Barrier Reef (12.0 g Si m^−2^; Maldonado et al. 2010). The sponge Si standing stock of the Bay of Brest is also similar to that found at bathyal depths at the westward slope of the Mauna Loa Volcano of Hawaii (12.6 g Si m^−2^; Maldonado et al. 2005), where a dense monospecific population of the highly-silicified hexactinellid *Sericolophus hawaiicus* occurs. Higher records of Si sponge stocks have only been reported from 1) the monospecific sponge ground of the hexactinellid *Vazella pourtalessii* at the Nova Scotian continental shelf, Canada (43.8 ± 74.6 g Si m^−2^; Maldonado et al. 2021), 2) the continental margins of Antarctica (178 g Si m^−2^; Gutt et al. 2013), where heavily silicified demosponges and hexactinellids co-occur, and 3) the singular epibathyal reefs of the hexactinellid *Aphrocallistes vastus* in British Columbia (Canada), where high densities of heavily silicified individuals grow on exposed skeletons of dead sponges, leading to outstanding accumulations of sponge silica in the form of siliceous reefs (4238 ± 924 g Si m^−2^; Chu et al. 2011). Altogether, these results support the idea that sponge communities are transient Si sinks (Maldonado et al. 2005, 2010) and suggest that relevant standing stocks of sponge silica are likely to occur not only in deep-sea and polar latitudes but also in shallow-water ecosystems from temperate latitudes, in which diverse and abundant sponge populations may easily develop (Van Soest et al. 2012; Maldonado et al. 2017).

The amount of sponge silica accumulated in only the superficial (upper centimeter) sediment layer of the Bay of Brest (1775 ± 162 × 10^3^ kg Si; Fig. 4) falls in the same order of magnitude than that accumulated in the living sponge fauna of the Bay (1215 ± 1876 × 10^3^ kg; Fig. 4). The Si stock in the superficial sediments is in the form of siliceous spicules that reach the sediments after sponge death (Chou et al. 2012; Lukowiak et al. 2013; Maldonado et al. 2019), the abundance of which appears to be proportional to the silica standing stock in the sponge communities living nearby (Bavestrello et al. 1996; Lukowiak et al. 2013). The amounts of sponge silica deposited on the superficial sediments of the Bay of Brest (0.799 – 2.505 mg Si g^−1^ sediment; Table 4) are among the highest determined in superficial sediments of continental margins from different oceans and seas (0.014 – 2.572 mg Si g^−1^ sediment; Sañé et al. 2013; Maldonado et al. 2019). For instance, the amount of sponge silica determined in the superficial sediments of the shallow muds (2.505 mg Si g^−1^ sediment, core SRQ1-IS05a; Table 4) is nearly identical to that determined in the superficial sediments of the slope of the Bransfield Strait in Antarctica (2.572 ± 0.861 mg Si g^−1^ sediment; Maldonado et al. 2019), where dense sponge aggregations occur (Ríos and Cristobo 2014; Kersken et al. 2016; Gutt et al. 2019). Combining the cores examined in this study (1.565 ± 0.241 mg Si g^−1^ sediment), the sediments of the Bay had about 60% more sponge silica than the average estimated for the sediments of continental margins in the global ocean (0.924 ± 0.854 mg Si g^−1^ sediment; Maldonado et al., 2019), indicating that sediments from areas where sponges abound become an important reservoir of biogenic silica.

### Silicon cycling through the sponge assemblage

The siliceous skeletons produced by the sponges as part of their annual growth are progressively accumulated within the sponge body for the lifespan, which is thought to range from years to decades or centuries in shallow-water sponge species from temperate latitudes (McMurray et al. 2008; Teixidó et al. 2011; McGrath et al. 2018). When a sponge dies or part of its body is removed or damaged by either natural or anthropogenic processes, the siliceous spicules within the sponge tissue are freed and end deposited on the superficial sediments (Chou et al. 2012; Lukowiak et al. 2013; Maldonado et al. 2019). In the Bay of Brest, the deposition of sponge silica was estimated to be 108 ± 9 × 10^3^ kg Si y^−1^ (Fig. 4). Such rate is about twice smaller than the rate at which sponges produce silica in the Bay (201 ± 372 × 10^3^ kg Si y^−1^; Fig. 4). That would mean that the sponge fauna of the Bay of Brest would be increasing at a rate of 92 ± 241 × 10^3^ kg Si y^−1^ (3.3 ± 8.6 × 10^6^ mol Si y^−1^), which would double the sponge Si standing stock (i.e., the sponge population) in about 13 years. In contrast, the sponge fauna survey of the Bay over the last 6 years and the long-term survey conducted in the area by the Marine Observatory of the IUEM since 1997 do not indicate that the sponge populations of the Bay are increasing at such high rate, rather the abundance and biomass of the sponge assemblages appear to be relatively constant in the long run (J. Grall, pers. comm.). Other reasons may also contribute to the imbalance between the annual silica production and deposition rates. First, the sponge mortality rate is unlikely to be constant every year. Indeed, a longer time frame (of at least a decade) is likely needed to capture the Si cycling dynamics in the sponge populations (McMurray et al. 2015; Bell et al. 2017). Second, there might be some partial rapid (<1y) dissolution of the spicules upon sponge death. Although sponge silica is more resistant to dissolution than that of others silicifiers (Rützler and Macintyre 1978; Erez et al. 1982; Maldonado et al. 2005, 2019), the most labile fraction of the sponge silica might be dissolved before the spicules being accumulated in the sediments. This would be in agreement with a recent study (Ng et al. 2020) that has measured through δ^30^ Si that the remineralization of silica from demosponges —the same Class of sponges as those occurring in the Bay of Brest— may be locally significant in superficial sediments where sponges abound. In the study, Ng and co-authors determined that the benthic Si effluxes from bottoms with sponges were from 2 to 10 times higher than those from sediments without sponges. Finally, it cannot be excluded that part of the sponge silica deposited on the Bay bottoms is exported out of the Bay during the monthly strong tidal currents during spring tides or during extreme storms events, the force of which has been suggested to partially transport the deposited sediments of the Bay (Beudin 2014). Further research is necessary to resolve which of these processes, or if a combination of them, could explain the imbalance between the annual sponge silica production and deposition rates at the Bay.

Through deposition, the sponge silica accumulates in the sediments of the Bay and is finally buried at an average rate of 0.06 cm y^−1^ (Gregoire et al. 2017; Ehrhold et al. 2021). Within the first centimeters of marine sediments, biogenic silica —no matter its origin— dissolves progressively until the interstitial water becomes saturated in dSi, an asymptotic condition in which biogenic silica dissolution typically ceases (Rickert et al. 2002; Sarmiento and Gruber 2006; Khalil et al. 2007). In the sediments of the Bay of Brest, the interstitial water reaches dSi asymptote at 10-15 cm burial depth (Raimonet et al. 2013), meaning that below that threshold, the amount of sponge silica buried annually (57 ± 18 × 10^3^ kg Si y^−1^; Fig. 4) is preserved definitively in the sediments of the Bay. The difference between the sponge silica deposited on superficial sediments and that buried definitively in the sediments (51 ± 18 × 10^3^ kg Si y^−1^; Fig. 4) is assumed to be dissolved as interstitial dSi during the early steps of burial. Such amount of dSi contributes to the saturation of interstitial water and ultimately feeds the dSi benthic efflux from the sediments of the Bay toward the water column, a recycling process that helps to sustain the populations of planktonic diatoms during the productive season in the Bay (Chauvaud et al. 2000; Ragueneau et al. 2002). Further investigation on the processes involved in the dissolution of sponge silica during the early steps of burial would help to quantify more accurately at which level the sponge Si contributes to the benthic dSi efflux.

### Sponge vs. diatom Si cycle at the Bay of Brest

The biogeochemical cycling of Si in the Bay of Brest through planktonic diatoms is well studied (e.g., Delmas and Tréguer, 1983; Del Amo et al., 1997; Chauvaud et al., 2000; Beucher et al., 2004) and summarized in Ragueneau et al. (2005). This budget was based on diatom Si flux rates and did not consider the sponge Si flux rates, nor the biogenic silica stocks of sponges and diatoms in the living communities and sediments.

To compare the contribution of diatoms and sponges to the Si budget of the Bay, the standing stocks and Si reservoirs in the form of diatom silica were determined from the literature. According to it, the standing stock of diatoms in the water column of the Bay ranges from 0.12 to 1.98 μmol Si L^−1^ over the year cycle (Beucher et al. 2004). Unlike sponges, diatom cells have an ephemeral life of only days. Therefore, the silica standing crop of planktonic diatoms is known to change drastically from week to week and over seasons, depending on nutrient availability and hydrological conditions (Sarthou et al. 2005; Falkowski and Oliver 2007; Armbrust 2009). If the annual mean of diatom silica in the Bay (0.9 μmol Si L^−1^; Beucher et al. 2004) is homogeneously extrapolated to the whole volume of the Bay of Brest (1.07 × 10^12^ L), an average diatom silica standing stock of 27 × 10^3^ kg Si (1.0 × 10^6^ mol Si) is obtained (Fig. 4). This figure integrates the seasonal variability occurring over the year cycle in the Bay, which range from a minimal value of 18 × 10^3^ kg Si (0.6 × 10^6^ mol Si) in winter to a maximum value of 36 × 10^3^ kg Si (1.3 × 10^6^ mol Si) in spring.

The fate of diatom silica in superficial sediments is largely influenced by the active filter-feeding of mollusks in the Bay, which defecate Si-rich feces that facilitate the retention of diatom silica at the sediment surface (Chauvaud et al. 2000; Ragueneau et al. 2002, 2005). The content of diatom silica in the superficial sediment of the Bay was determined from the analysis of 86 sediment samples from 43 stations across the Bay, which capture both the intra-annual variability (43 samples were sampled in winter and 43 in late summer) and the different depositional environments of the Bay (Song and Ragueneau 2007). It resulted in an average diatom silica reservoir in the superficial sediments of the Bay of 248 × 10^3^ kg Si (8.8 × 10^6^ mol Si; Fig. 4).

When the Si stocks and fluxes of sponges are compared with those of planktonic diatoms, there are marked differences (Fig. 4). The rates at which Si is processed through diatoms are one order of magnitude higher than those through sponges. On the contrary, diatom Si stocks are between 7 to 45 times smaller than those of sponges (Fig. 4), in agreement with the only study to date comparing Si standing stocks in sponges and diatoms in a coastal ecosystem (Maldonado et al. 2010). These differences indicate that sponges and diatoms play their respective roles in the Si budget of the Bay at different speeds and through different mechanisms. The turnover of the diatom silica standing stock (i.e., standing stock: production rate) in the Bay is about 3 days, whereas that of sponge silica is about 6 years, that is, approximately 800 times slower than that of diatom silica. Similarly, the turnover of diatom silica stock in superficial sediments (i.e., reservoir in sediment: deposition rate) is about 1 month, whereas that of sponge silica is about 16 years (ca. 170 times that of diatom silica). This means that the cycling of Si through diatoms is comparatively faster, with Si mainly cycling through them (repeatedly over a year) rather than being accumulated. In contrast, Si accumulates in huge amounts in sponges over long periods (≥10 y), being slowly processed through them. These results show that impacts on either the sponge populations or the sediments nearby sponge aggregations could have a long-term impact on the sponge Si cycling dynamics, which will require decades to be restored.

## Conclusion

In shallow-water ecosystems, planktonic and benthic silicifiers have to share the dSi pool and compete to incorporate this nutrient, which is critical for the growth of both organisms. Our study highlights that even in coastal, shallow-water systems with a high primary productivity dominated by planktonic diatoms —such as the Bay of Brest, sponges may account for a stock of biogenic silica as large as 89-98%. This stock cycles slowly compared to that of diatoms and accumulates bSi in the sediments. These results suggest that the Si cycling through sponge communities is substantially different from that through diatom assemblages. Therefore, the comparison between the roles of these two groups of silicifiers is not straightforward due to their contrasting biological features (e.g., benthic and long living vs planktonic and short living). Yet these two types of silicifiers need to be quantitatively integrated in future approaches, if we aim to understand in depth the intricacies of the Si cycle in coastal systems.

## Acknowledgments

The authors thank Erwan Amice, Isabelle Bihannic, and Thierry Le Bec for assistance during underwater fieldwork and for pictures of Fig. 1b-g. Marta García Puig and Marcos Navas Durán are thanked for helping with the data management of spicule counting. The authors also thank the staff maintaining the Lanvéoc database for making public their information on nutrient availability at the bottom waters of the Bay of Brest and Sébastien Hervé for the artwork of Fig. 4. Jill Sutton and Fiz Fernández are especially thanked for their comments on the manuscript. This research was supported by two Spanish Ministry grants (CTM2015-67221-R and MICIU: #PID2019-108627RB-I00) to MM, the French National research program EC2CO (grant 12735 – AO2020) to JG, and the ISblue project, Interdisciplinary graduate school for the blue planet (ANR-17-EURE-0015), co-funded by a grant from the French government under the program “Investissements d’Avenir”, to MLA. MLA thanks the Xunta de Galicia for her postdoctoral grant (IN606B-2019/002), which also supported this work. The authors declare no conflict of interest.

## Author Contributions

MLA and MM conceived and designed the study. MLA conducted the sponge fieldwork, with the help of JG and AL. MLA, MM, and CS taxonomically identified the sponges. MLA processed the sponge samples to determine specific silica content. AH sampled the sediment cores and provided sedimentary data. JG and AH provided the bottom mapping data and MLA developed the benthic habitat and spicule accumulation layers. MLA, CG, and CS conducted sponge silica determination under the microscope. MLA analyzed and interpreted the data and drafted the manuscript, with invaluable inputs made by MM and AL. All authors contributed with comments to the manuscript and approved the submitted version.

## Supplemental Information

**Supplementary Table 1.**
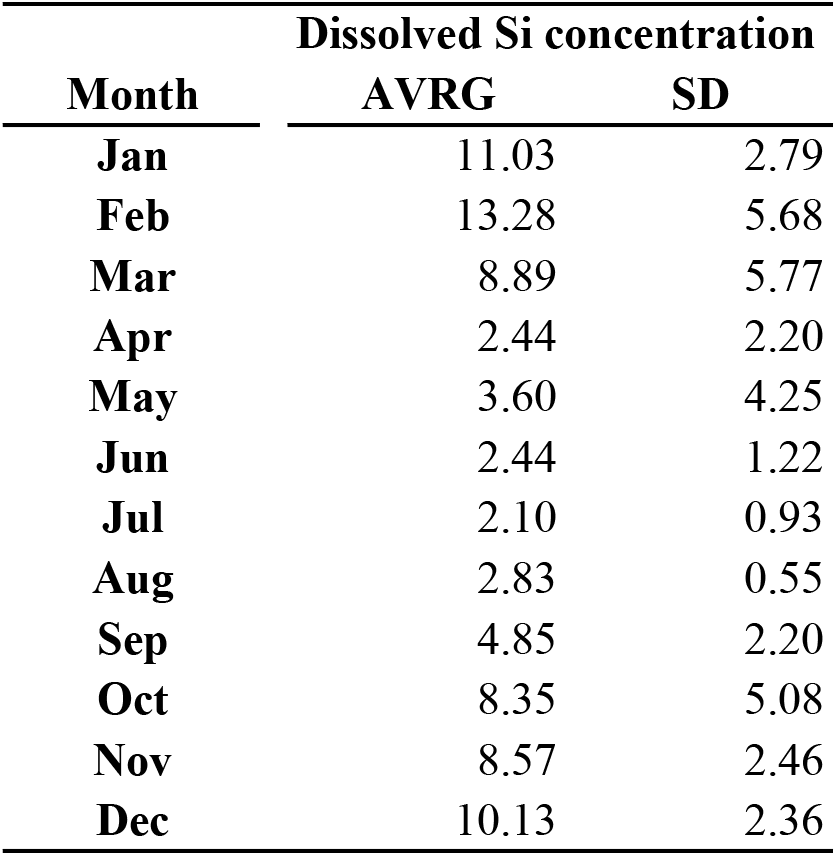
Average concentration (±SD) of dissolved silicon (μmol Si L-1) at the bottom waters of the Bay of Brest. Data come from the Lanvéoc long-term survey series (48.295777° N, 4.454758° W), which sampled bottom water with a 1L syringe manually by scientific divers just over seabed every two weeks at high tide. Data integrated dissolved silicon concentrations from the last decade (2012-2021).

**Supplementary Table 2.**
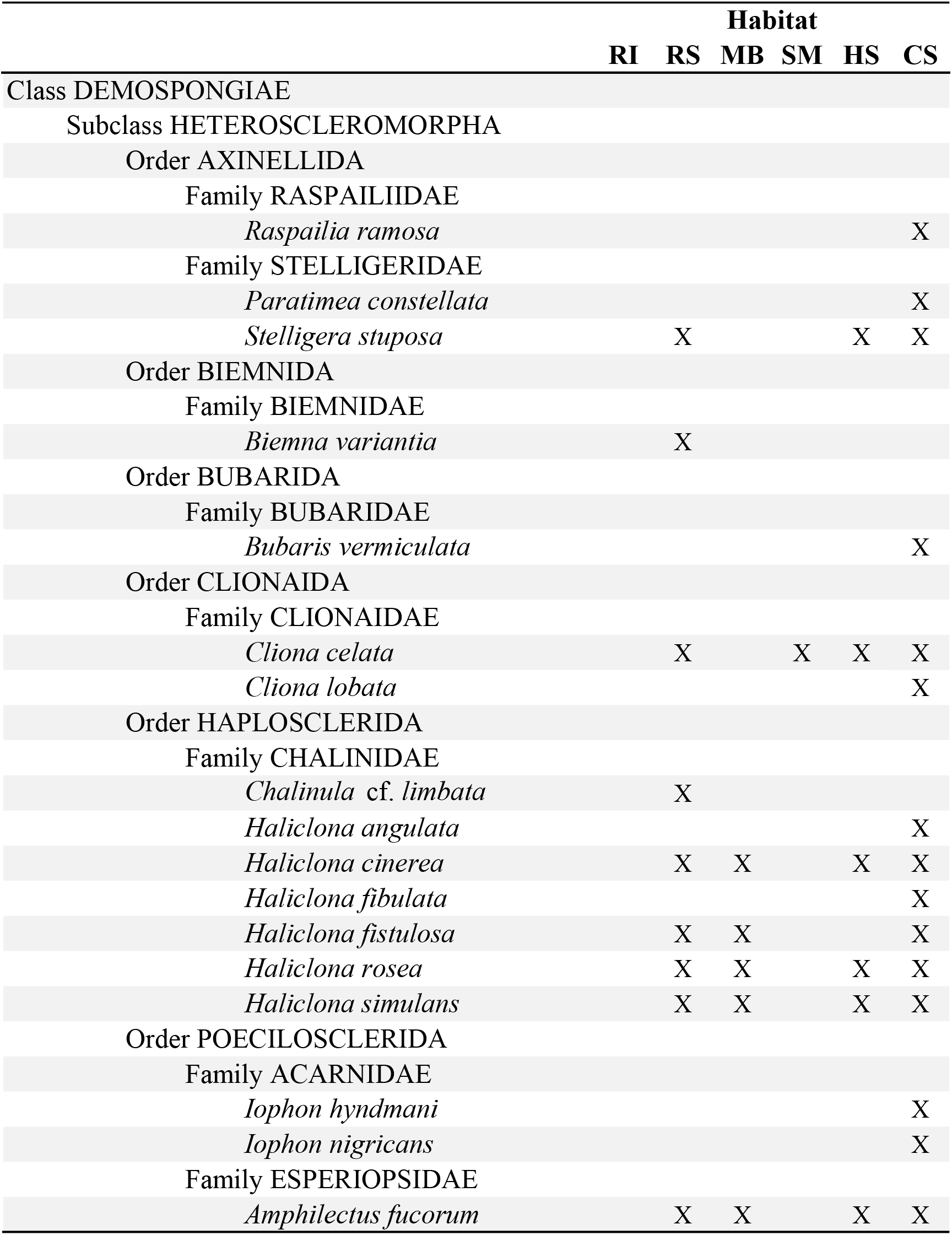
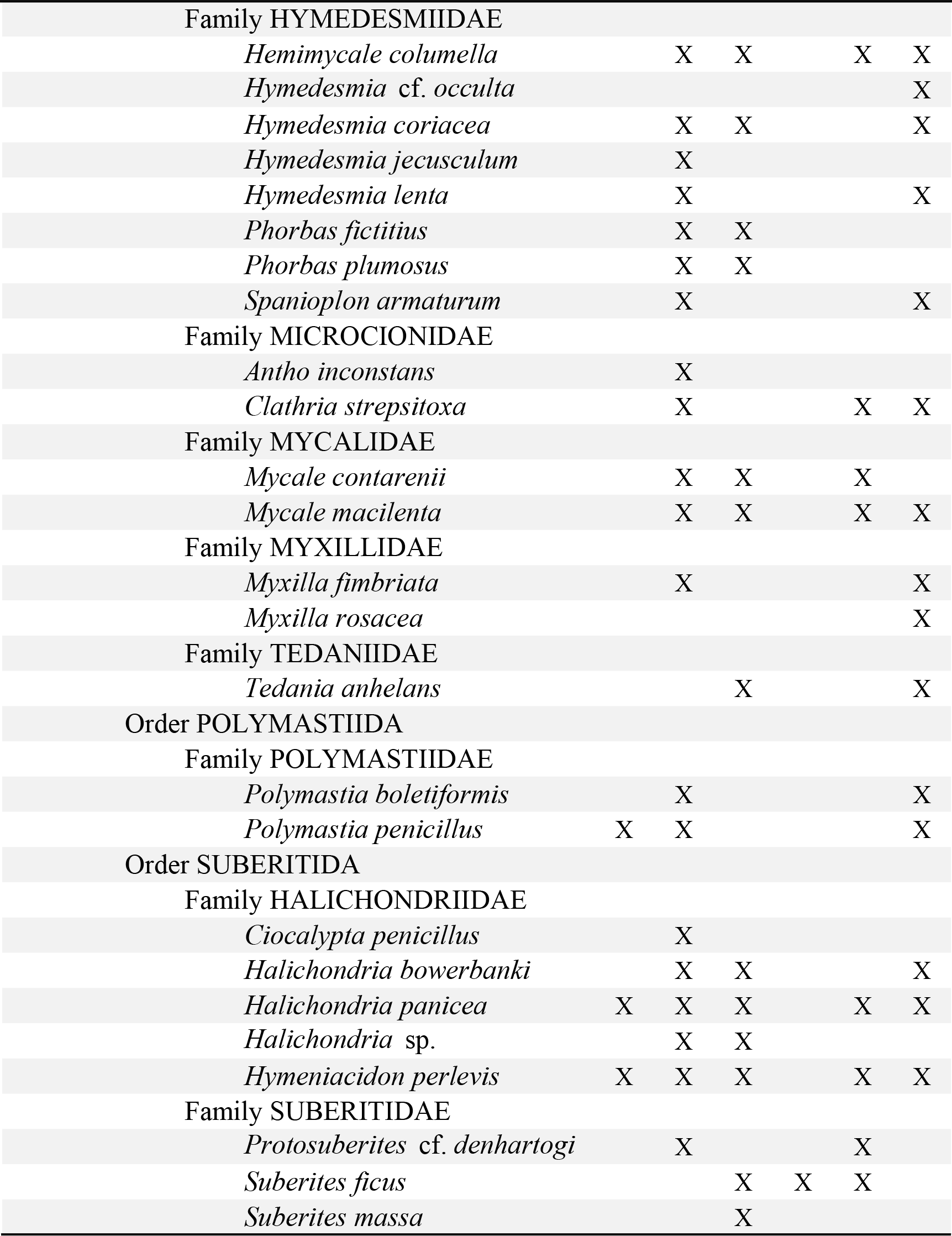
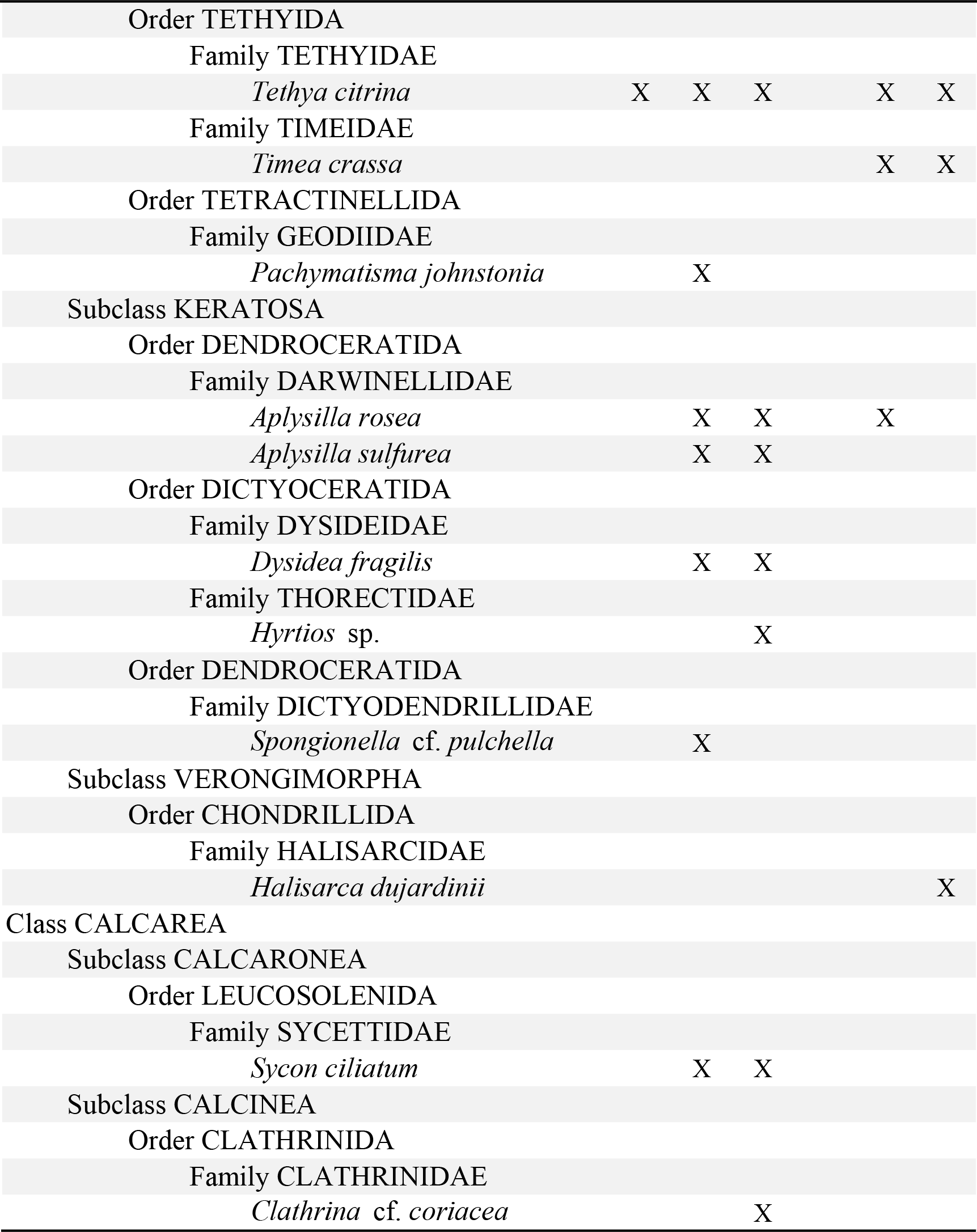
List of the sponge species of the Bay of Brest (France). Presence at each habitat is indicated. RI, rocky intertidal; RS, rocky subtidal; MB, maerl beds; SM, shallow mud; HS, heterogeneous sediments; CS, circalittoral coarse sediments.

## References

Armbrust, E. V. 2009. The life of diatoms in the world’s oceans. Nature 459: 185–192. doi:10.1038/nature08057

Ávila, E., and R. Riosmena-Rodriguez. 2011. A preliminary evaluation of shallow-water rhodolith beds in Bahia Magdalena, Mexico. Braz. J. Oceanogr. 59: 365–375. doi:10.1590/S1679-87592011000400007

Bavestrello, G., R. Cattaneo-Vietti, C. Cerrano, S. Cerutti, and M. Sará. 1996. Contribution of sponge spicules to the composition of biogenic silica in the Ligurian Sea. Mar. Ecol. 17: 41–50. doi:10.1111/j.1439-0485.1996.tb00488.x

Bell, J. J., A. Biggerstaff, T. Bates, H. Bennett, J. Marlow, E. McGrath, and M. Shaffer. 2017. Sponge monitoring: moving beyond diversity and abundance measures. Ecol. Indic. 78: 470–488. doi:10.1016/j.ecolind.2017.03.001

Beucher, C., P. Treguer, R. Corvaisier, A. M. Hapette, and M. Elskens. 2004. Production and dissolution of biosilica, and changing microphytoplankton dominance in the Bay of Brest (France). Mar. Ecol. Prog. Ser. 267: 57–69. doi:10.3354/meps267057

Beudin, A. 2014. Dynamique et échanges sédimentaires en rade de Brest impactés par l’invasion de crépidules. Ph.D. thesis. Université de Bretagne Occidentale.

Chauvaud, L., F. Jean, O. Ragueneau, and G. Thouzeau. 2000. Long-term variation of the Bay of Brest ecosystem: benthic-pelagic coupling revisited. Mar. Ecol. Prog. Ser. 200: 35–48. doi:10.3354/meps200035

Chou, Y., J. Y. Lou, C.-T. A. Chen, and L.-L. Liu. 2012. Spatial distribution of sponge spicules in sediments around Taiwan and the Sunda Shelf. J. Oceanogr. 68: 905–912. doi:10.1007/s10872-012-0143-7

Chu, J. W. F., M. Maldonado, G. Yahel, and S. P. Leys. 2011. Glass sponge reefs as a silicon sink. Mar. Ecol. Prog. Ser. 441: 1–14. doi:10.3354/meps09381

Davidson, K., R. J. Gowen, P. J. Harrison, L. E. Fleming, P. Hoagland, and G. Moschonas. 2014. Anthropogenic nutrients and harmful algae in coastal waters. J. Environ. Manage. 146: 206–216. doi:10.1016/j.jenvman.2014.07.002

Del Amo, Y., B. Quéguiner, P. Tréguer, H. Breton, and L. Lampert. 1997. Impacts of high-nitrate freshwater inputs on macrotidal ecosystems. II. Specific role of the silicic acid pump in the year-round dominance of diatoms in the Bay of Brest (France). Mar. Ecol. Prog. Ser. 161: 225–237. doi:10.3354/meps161225

Delmas, R., and P. Tréguer. 1983. Evolution saisonnière des nutriments dans un écosystème eutrophe de l’Europe Occidentale (la Rade de Brest). Interactions marines et terrestres. Oceanol. Acta 6: 345–356.

Delmas, R., and P. Tréguer. 1985. Simulation de l’évolution de paramètres physiques, chimiques, et de la biomasse phytoplanctonlque en période printanière dans un écosystème littoral macrotidal. Oceanis 11: 197–211.

DeMaster, D. J. 2003. The diagenesis of biogenic silica: chemical transformations occurring in the water column, seabed, and crust, p. 87–98. *In* F.T. Mackenzie [ed.], Treatise on Geochemistry. Elsevier.

Derrien-Courtel, S., T. Androuin, E. Ar Gall, and others. 2019. Surveillance du Benthos du littoral Breton. Années 2017-2018 Rapport final. Museum d’Histoire Naturelle de Paris.

Eberhardt, L. L. 1978. Appraising variability in population studies. J. Wildlife Manage. 42: 207. doi:10.2307/3800260

Ehrhold, A., G. Gregoire, S. Sabine, G. Jouet, and P. Le Roy. 2016. Present-day sedimentation rates and evolution since the last maximum flooding surface event in the Bay of Brest (W-N France). Proceedings of the American Geophysical Union, AGU, Fall Meeting 2016.

Ehrhold, A., G. Jouet, P. Le Roy, and others. 2021. Fossil maerl beds as coastal indicators of late Holocene palaeo-environmental evolution in the Bay of Brest (Western France). Palaeogeography, Palaeoclimatology, Palaeoecology 577: 110525. doi:10.1016/j.palaeo.2021.110525

Erez, J., K. Takahashi, and S. Honjo. 1982. In-situ dissolution experiment of radiolaria in the central North Pacific ocean. Earth Planet. Sci. Lett. 59: 245–254. doi:10.1016/0012-821X(82)90129-7

Falkowski, P. G., and M. J. Oliver. 2007. Mix and match: how climate selects phytoplankton. Nat. Rev. Microbiol. 5: 813–819. doi:10.1038/nrmicro1751

Glibert, P. M., and M. A. Burford. 2017. Globally changing nutrient loads and harmful algal blooms. Recent advances, new paradigms, and continuing challenges. Oceanography 30: 58–69.

Grall, J., and M. Glémarec. 1997. Biodiversity of maerl beds in Brittany: functional approach and anthropogenic impacts. Vie et Milieu 47: 339–349.

Gregoire, G., A. Ehrhold, P. Le Roy, G. Jouet, and T. Garlan. 2016. Modern morpho-sedimentological patterns in a tide-dominated estuary system: the Bay of Brest (west Britanny, France). J. Maps 12: 1152–1159. doi:10.1080/17445647.2016.1139514

Gregoire, G., P. Le Roy, A. Ehrhold, G. Jouet, and T. Garlan. 2017. Control factors of Holocene sedimentary infilling in a semi-closed tidal estuarine-like system: the bay of Brest (France). Mar. Geol. 385: 84–100. doi:10.1016/j.margeo.2016.11.005

Gutt, J., J. Arndt, C. Kraan, B. Dorschel, M. Schröder, A. Bracher, and D. Piepenburg. 2019. Benthic communities and their drivers: a spatial analysis off the Antarctic Peninsula. Limnol. Oceanogr. 64: 2341–2357. doi:10.1002/lno.11187

Gutt, J., A. Bohmer, and W. Dimmler. 2013. Antarctic sponge spicule mats shape macrobenthic diversity and act as a silicon trap. Mar. Ecol. Prog. Ser. 480: 57–71. doi:10.3354/meps10226

Hily, C., P. Potin, and J.-Y. Floch. 1992. Structure of subtidal algal assemblages on soft-bottom sediments: fauna/flora interactions and role of disturbances in the Bay of Brest, France. Mar. Ecol. Prog. Ser. 85: 115–130. doi:10.3354/meps085115

Hooper, J. N. A. 2019. Sponges, p. 170–186. *In* P.A. Hutchings, M. Kingsford, and O. Hoegh-Guldberg [eds.], The Great Barrier Reef: biology, environment and management.

Jean, F. 1994. Modélisation à l’état stable des transferts de carbone dans le réseau trophique benthique de la rade de Brest (France). Ph.D. thesis. Université de Bretagne Occidentale.

Kersken, D., B. Feldmeyer, and D. Janussen. 2016. Sponge communities of the Antarctic Peninsula: influence of environmental variables on species composition and richness. Polar Biol. 39: 851–862. doi:10.1007/s00300-015-1875-9

Khalil, K., C. Rabouille, M. Gallinari, K. Soetaert, D. J. DeMaster, and O. Ragueneau. 2007. Constraining biogenic silica dissolution in marine sediments: a comparison between diagenetic models and experimental dissolution rates. Mar. Chem. 106: 223–238. doi:10.1016/j.marchem.2006.12.004

Kristiansen, S., and E. E. Hoell. 2002. The importance of silicon for marine production. Hydrobiologia 484: 21–31. doi:10.1023/A:1021392618824

Lambert, C., M. Vidal, A. Penaud, N. Combourieu-Nebout, V. Lebreton, O. Ragueneau, and G. Gregoire. 2017. Modern palynological record in the Bay of Brest (NW France): Signal calibration for palaeo-reconstructions. Review of Palaeobotany and Palynology 244: 13–25. doi:10.1016/j.revpalbo.2017.04.005

Le Pape, O., Y. Del Amo, A. Menesguen, A. Aminot, B. Quéquiner, and P. Tréguer. 1996. Resistance of a coastal ecosystem to increasing eutrophic conditions: the Bay of Brest (France), a semi-enclosed zone of Western Europe. Cont. Shelf Res. 16: 1885–1907. doi:10.1016/0278-4343(95)00068-2

López-Acosta, M., A. Leynaert, J. Grall, and M. Maldonado. 2018. Silicon consumption kinetics by marine sponges: an assessment of their role at the ecosystem level. Limnol. Oceanogr. 63: 2508–2522. doi:10.1002/lno.10956

López-Acosta, M., A. Leynaert, and M. Maldonado. 2016. Silicon consumption in two shallow-water sponges with contrasting biological features. Limnol. Oceanogr. 61: 2139–2150. doi:10.1002/lno.10359

Lukowiak, M., A. Pisera, and A. O’Dea. 2013. Do spicules in sediments reflect the living sponge community? A test in a Caribbean shallow-water lagoon. PALAIOS 28: 373–385. doi:10.2110/palo.2012.p12-082r

Maldonado, M., R. Aguilar, R. J. Bannister, and others. 2017. Sponge grounds as key marine habitats: a synthetic review of types, structure, functional roles, and conservation concerns, p. 145–184. *In* S. Rossi, L. Bramanti, A. Gori, and C. Orejas Saco del Valle [eds.], Marine Animal Forests: The Ecology of Benthic Biodiversity Hotspots. Springer International Publishing.

Maldonado, M., L. Beazley, M. López-Acosta, E. Kenchington, B. Casault, U. Hanz, and F. Mienis. 2021. Massive silicon utilization facilitated by a benthic-pelagic coupled feedback sustains deep-sea sponge aggregations. Limnol Oceanogr 66: 366–391. doi:10.1002/lno.11610

Maldonado, M., C. Carmona, Z. Velásquez, A. Puig, A. Cruzado, A. López, and C. M. Young. 2005. Siliceous sponges as a silicon sink: an overlooked aspect of benthopelagic coupling in the marine silicon cycle. Limnol. Oceanogr. 50: 799–809. doi:10.4319/lo.2005.50.3.0799

Maldonado, M., M. López-Acosta, L. Beazley, E. Kenchington, V. Koutsouveli, and A. Riesgo. 2020. Cooperation between passive and active silicon transporters clarifies the ecophysiology and evolution of biosilicification in sponges. Sci. Adv. 6: eaba9322. doi:10.1126/sciadv.aba9322

Maldonado, M., M. López-Acosta, C. Sitjà, M. García-Puig, C. Galobart, G. Ercilla, and A. Leynaert. 2019. Sponge skeletons as an important sink of silicon in the global oceans. Nat. Geosci. 12: 815–822. doi:10.1038/s41561-019-0430-7

Maldonado, M., L. Navarro, A. Grasa, A. González, and I. Vaquerizo. 2011. Silicon uptake by sponges: a twist to understanding nutrient cycling on continental margins. Sci Rep 1: 8. doi:10.1038/srep00030

Maldonado, M., A. Riesgo, A. Bucci, and K. Rützler. 2010. Revisiting silicon budgets at a tropical continental shelf: silica standing stocks in sponges surpass those in diatoms. Limnol. Oceanogr. 55: 2001–2010. doi:10.4319/lo.2010.55.5.2001

Malviya, S., E. Scalco, S. Audic, and others. 2016. Insights into global diatom distribution and diversity in the world’s ocean. Proc Natl Acad Sci USA 113: E1516–E1525. doi:10.1073/pnas.1509523113

McGrath, E. C., L. Woods, J. Jompa, A. Haris, and J. J. Bell. 2018. Growth and longevity in giant barrel sponges: redwoods of the reef or pines in the Indo-Pacific? Sci Rep 8: 15317. doi:10.1038/s41598-018-33294-1

McMurray, S. E., J. E. Blum, and J. R. Pawlik. 2008. Redwood of the reef: growth and age of the giant barrel sponge *Xestospongia muta* in the Florida Keys. Mar. Biol. 155: 159–171. doi:10.1007/s00227-008-1014-z

McMurray, S. E., C. M. Finelli, and J. R. Pawlik. 2015. Population dynamics of giant barrel sponges on Florida coral reefs. J. Exp. Mar. Biol. Ecol. 473: 73–80. doi:10.1016/j.jembe.2015.08.007

Neill, K. F., W. A. Nelson, R. D’Archino, D. Leduc, and T. J. Farr. 2015. Northern New Zealand rhodoliths: assessing faunal and floral diversity in physically contrasting beds. Mar Biodiv 45: 63–75. doi:10.1007/s12526-014-0229-0

Ng, H. C., L. Cassarino, R. A. Pickering, E. M. S. Woodward, S. J. Hammond, and K. R. Hendry. 2020. Sediment efflux of silicon on the Greenland margin and implications for the marine silicon cycle. Earth Planet. Sci. Lett. 529: 115877. doi:10.1016/j.epsl.2019.115877

QGIS Development Team. 2020. QGIS Geographic Information System, Open Source Geospatial Foundation Project.

Quéguiner, B., and P. Tréguer. 1984. Studies on the phytoplankton in the bay of Brest (western Europe). Seasonal variations in composition, biomass and production in relation to hydrological and chemical features (1981-1982). Bot. Mar. 27: 449–459. doi:10.1515/botm.1984.27.10.449

Ragueneau, O., L. Chauvaud, A. Leynaert, and others. 2002. Direct evidence of a biologically active coastal silicate pump: ecological implications. Limnol. Oceanogr. 47: 1849–1854. doi:10.4319/lo.2002.47.6.1849

Ragueneau, O., L. Chauvaud, B. Moriceau, A. Leynaert, G. Thouzeau, A. Donval, F. Le Loc’h, and F. Jean. 2005. Biodeposition by an invasive suspension feeder impacts the biogeochemical cycle of Si in a coastal ecosystem (Bay of Brest, France). Biogeochemistry 75: 19–41. doi:10.1007/s10533-004-5677-3

Ragueneau, O., D. J. Conley, A. Leynaert, S. N. Longphuirt, and C. P. Slomp. 2006. Role of diatoms in silica cycling and coastal marine food webs, p. 163–195. *In* V. Ittekkot, D. Unger, C. Humborg, and N.T. An [eds.], The silicon cycle: human perturbations and impacts on aquatic systems. Island Press.

Ragueneau, O., M. Raimonet, C. Mazé, and others. 2018. The impossible sustainability of the Bay of Brest? Fifty years of ecosystem changes, interdisciplinary knowledge construction and key questions at the science-policy-community interface. Front. Mar. Sci. 5: 124. doi:10.3389/fmars.2018.00124

Raimonet, M., O. Ragueneau, F. Andrieux-Loyer, X. Philippon, R. Kerouel, M. Le Goff, and L. Mémery. 2013. Spatio-temporal variability in benthic silica cycling in two macrotidal estuaries: Causes and consequences for local to global studies. Estuar. Coast. Shelf Sci. 119: 31–43. doi:10.1016/j.ecss.2012.12.008

Reincke, T., and D. Barthel. 1997. Silica uptake kinetics of *Halichondria panicea* in Kiel Bight. Marine Biology 129: 591–593. doi:10.1007/s002270050200

Rickert, D., M. Schlüter, and K. Wallmann. 2002. Dissolution kinetics of biogenic silica from the water column to the sediments. Geochim. Cosmochim. Acta 66: 439–455. doi:10.1016/S0016-7037(01)00757-8

Ríos, P., and J. Cristobo. 2014. Antarctic Porifera database from the Spanish benthic expeditions. ZooKeys 401: 1–10. doi:10.3897/zookeys.401.5522

Rützler, K., and I. G. Macintyre. 1978. Siliceous sponge spicules in coral-reef sediments. Mar. Biol. 49: 147–159. doi:10.1007/bf00387114

Salomon, J. C., and M. Breton. 1991. Numerical study of the dispersive capacity of the Bay of Brest, France, towards dissolved substances, p. 459–464. *In* Environmental hydraulics.

Sandford, F. 2003. Physical and chemical analysis of the siliceous skeletons in six sponges of two groups (Demospongiae and Hexactinellida). Microsc. Res. Tech. 62: 336–355. doi:10.1002/jemt.10400

Sañé, E., E. Isla, M. Á. Bárcena, and D. J. DeMaster. 2013. A shift in the biogenic silica of sediment in the Larsen B continental shelf, off the Eastern Antarctic Peninsula, resulting from climate change V. Magar [ed.]. PLoS ONE 8: e52632. doi:10.1371/journal.pone.0052632

Sarmiento, J., and N. Gruber. 2006. Ocean biogeochemical dynamics, Princeton University Press.

Sarthou, G., K. R. Timmermans, S. Blain, and P. Tréguer. 2005. Growth physiology and fate of diatoms in the ocean: a review. J. Sea Res. 53: 25–42. doi:10.1016/j.seares.2004.01.007

Sciberras, M., M. Rizzo, J. R. Mifsud, K. Camilleri, J. A. Borg, E. Lanfranco, and P. J. Schembri. 2009. Habitat structure and biological characteristics of a maerl bed off the northeastern coast of the Maltese Islands (central Mediterranean). Mar. Biodiv. 39: 251–264. doi:10.1007/s12526-009-0017-4

Song, Y. P., and O. Ragueneau. 2007. Le dosage de bSiO2 dans le sédiment de la rade de Brest. Master thesis. Université de Bretagne Occidentale.

Teixidó, N., J. Garrabou, and J.-G. Harmelin. 2011. Low dynamics, high longevity and persistence of sessile structural species dwelling on Mediterranean coralligenous outcrops S. Thrush [ed.]. PLoS ONE 6: e23744. doi:10.1371/journal.pone.0023744

Thorel, M., P. Claquin, M. Schapira, and others. 2017. Nutrient ratios influence variability in *Pseudo-nitzschia* species diversity and particulate domoic acid production in the Bay of Seine (France). Harmful Algae 68: 192–205. doi:10.1016/j.hal.2017.07.005

Tréguer, P. J., J. N. Sutton, M. Brzezinski, and others. 2021. Reviews and syntheses: The biogeochemical cycle of silicon in the modern ocean. Biogeosciences 18: 1269–1289. doi:10.5194/bg-18-1269-2021

Van Soest, R. W. M., N. Boury-Esnault, J. Vacelet, and others. 2012. Global diversity of sponges (Porifera). PLOS ONE 7: e35105. doi:10.1371/journal.pone.0035105

